# Short internal open reading frames regulate the translation of N-terminally truncated proteoforms

**DOI:** 10.1101/2023.11.09.566418

**Authors:** Raphael Fettig, Zita Gonda, Niklas Walter, Paul Sallmann, Christiane Thanisch, Markus Winter, Susanne Bauer, Lei Zhang, Greta Linden, Margarethe Litfin, Christelle Etard, Olivier Armant, Olalla Vázquez, Olivier Kassel

**Affiliations:** Karlsruhe Institute of Technology (KIT), Institute for Biological and Chemical Systems - Biological Information Processing (IBCS-BIP), Karlsruhe, Germany; Philipps-Universität Marburg, Faculty of Chemistry, Marburg, Germany; Institut de Radioprotection et de Sûreté Nucléaire (IRSN), PSE-ENV/SERPEN/LECO, Cadarache, France; Philipps-Universität Marburg, Center for Synthetic Microbiology (SYNMIKRO), Marburg, Germany

**Keywords:** myogenesis/proteoforms/short ORFs/translation/Trip6

## Abstract

Internal translation initiation sites, as revealed by ribosome profiling experiments can potentially drive the translation of many N-terminally truncated proteoforms. We have identified an mRNA cis-regulatory element which regulates their translation. nTRIP6 represents a short nuclear proteoform of the cytoplasmic protein TRIP6. We have previously reported that nTRIP6 regulates the dynamics of skeletal muscle progenitor differentiation. Here we show that nTRIP6 is generated by translation initiation at an internal AUG. Its translation is repressed by an internal short open reading frame (sORF) immediately upstream of the nTRIP6 AUG. Consistent with this representing a more general regulatory feature, we have identified other N-terminally truncated proteoforms which are repressed by internal sORFs. In an *in vitro* model of myogenic differentiation, the translation of nTRIP6 is transiently de-repressed in a mechanistic Target of Rapamycin Complex 1-dependent manner. Thus, the translation of N-terminally truncated proteoforms can be regulated independently of the canonical ORF.

## Introduction

The number of proteins that can be expressed in a cell far exceeds the number of genes. Alternative splicing and alternative translation of mRNAs drastically increase the coding capacity of the genome by generating multiple proteoforms. How the expression of these various proteoforms is regulated in a physiological context is still poorly understood, in particular in the case of alternative translation. In some cases, the proteoforms regulate the function of the “canonical” proteins (Kelemen *et al*, 2013; Orre *et al*, 2019; Bogaert *et al*, 2020), as for example the case of truncated transcription factors (TFs) which act as dominant-negative inhibitors.

However, there are also examples of proteoforms which exert a function distinct from that of their canonical counterparts. This occurs in particular when the canonical protein and its variant are located in different subcellular compartment s(Kelemen *et al*, 2013; Orre *et al*, 2019; Bogaert *et al*, 2020). The protein TRIP6 (Lin & Lin, 2011; Willier *et al*, 2011) represents one such example. TRIP6 belongs to the ZYXIN family of LIM domain-containing proteins that are localized in the cytosol and enriched at sites of cell adhesion where they regulate adhesion and migration. These proteins are also present in the nucleus where they exert transcriptional co-regulator functions for various TFs (Hervy *et al*, 2006). In the case of TRIP6, this nuclear function is exerted by a shorter, exclusively nuclear proteoform, which we have termed nTRIP6 (Kassel *et al*, 2004; Diefenbacher *et al*, 2008, 2010; Kemler *et al*, 2016). We have recently reported a role for nTRIP6 in the regulation of myogenesis (Norizadeh Abbariki *et al*, 2021). This process relies on resident adult stem cells, the so-called satellite cells (Schmidt *et al*, 2019). Upon muscle damage, these cells are activated and give rise to proliferating progenitors, the myoblasts. These then differentiate into committed precursors, the myocytes, which finally fuse together to form multinucleated myofibres. We have shown in an *in vitro* model of myogenesis that nTRIP6 prevents premature differentiation of myoblasts into myocytes. This repression is required for the subsequent late differentiation and fusion of myocytes. Furthermore, the expression of nTRIP6 is transiently upregulated at the transition between proliferation and differentiation, while that of TRIP6 is not (Norizadeh Abbariki *et al*, 2021).

Here, we report that nTRIP6 is expressed by alternative translation initiation at an internal AUG and that a short open reading frame (ORF) within the *Trip6* coding sequence represses nTRIP6 translation. Furthermore, the mechanistic target of rapamycin complex 1 (mTORC1) transiently de-represses nTRIP6 translation during early myogenesis. Based on this prototypical example, we identified other mRNAs in which short internal ORFs regulate the translation of N-terminally truncated proteoforms. Thus, our work reveals novel translation cis-regulatory elements that we have termed internal upstream ORFs (iuORFs).

## Results and discussion

### nTRIP6 is generated by alternative translation at an internal AUG

A nuclear export signal (NES) within the N-terminal pre-LIM region of TRIP6 is responsible for its cytosolic localization (Lin & Lin, 2011; Willier *et al*, 2011). We therefore predicted that the nuclear proteoform nTRIP6 lacks a functional NES. However, it should still harbor the C-terminal LIM domains as well as the interaction domains within the pre-LIM region which are required for its transcriptional co-regulator function (Kassel *et al*, 2004; Diefenbacher *et al*, 2014; Kemler *et al*, 2016). Transfection of NIH-3T3 fibroblasts with a C-terminally tagged *Trip6* construct gave rise to two proteins with sizes corresponding to those of TRIP6 and nTRIP6 (Fig 1A,B). Deletion of the annotated translation initiation codon (AUG1) abolished the expression of the long proteoform but strongly increased the expression of the short proteoform (Fig 1A,B). This result strongly suggests that nTRIP6 arises from alternative translation initiation after leaky scanning, the process by which the scanning ribosome skips the suboptimal first initiation codon and initiates at a downstream site (Kozak, 2002). Indeed, the *Trip6* AUG1 is immediately followed by a conserved T and not by the optimal G. Still, TRIP6 is expressed at very high levels compared to nTRIP6 in all cells tested (Kassel *et al*, 2004; Diefenbacher *et al*, 2008, 2010; Kemler *et al*, 2016). This difference in the relative levels of the two proteoforms is compatible with their functions. Indeed, TRIP6 is a regulator of the cytoskeleton whereas nTRIP6 is a transcriptional regulator. These are generally expressed at significant lower levels than cytoskeletal regulatory proteins (Beck *et al*, 2011). We identified a putative initiation codon (AUG2), in frame with AUG1 and located in the middle of the NES-encoding sequence. Mutation of AUG2 into a non-initiating codon abrogated the expression of the short proteoform (Fig 1A,B). To confirm that nTRIP6 is translated at AUG2, we designed a peptide nucleic acid (PNA) to block translation initiation (Marin *et al*, 2004; Gupta *et al*, 2017) at this site (Figure EV1). In an *in vitro* transcription/translation assay, the AUG2-targeting PNA inhibited the translation of the short proteoform, as compared to a mispaired control PNA (Misp), without affecting that of the long proteoform (Fig 1C). Deletion of AUG1 abolished the translation of the long proteoform and increased the translation of the short proteoform, which was inhibited by the AUG2 PNA (Fig 1C). Thus, nTRIP6 arises from translation initiation at AUG2 after leaky scanning. Given that AUG2 is located in the middle of the NES-encoding sequence and that the NES is the main determinant of the cytosolic localization of TRIP6 (Lin & Lin, 2011; Willier *et al*, 2011), the nuclear localization of nTRIP6 (Kassel *et al*, 2004; Kemler *et al*, 2016) is a logical consequence of the truncated NES.

**Figure 1:**
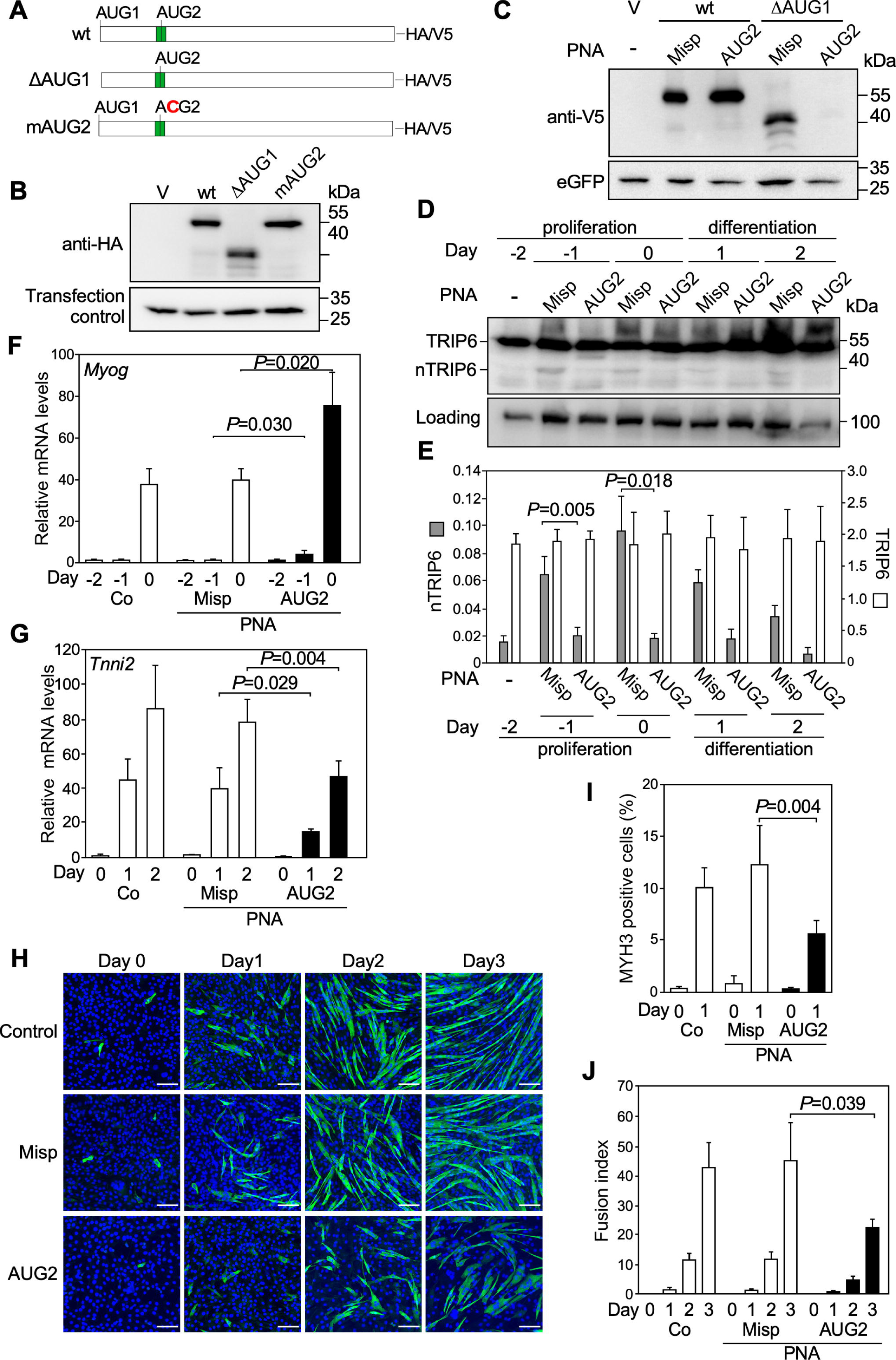
nTRIP6 is generated by alternative translation at an internal AUG. A Schematic representation of the constructs used. The green box represents the Nuclear Export Signal encoding sequence. B Lysates of NIH-3T3 fibroblasts transfected with either an empty vector (V) or the indicated HA-tagged TRIP6 constructs, together with eGFP as a transfection control were subjected to Western blotting using antibodies against the HA tag and eGFP. Results are representative of three independent experiments. C *In vitro* transcription and translation of V5-tagged Trip6 CDS constructs with (wt) or without the first AUG (ΔAUG1), as well as of eGFP as in internal translation control, in the presence of a peptide nucleic acid (PNA) targeting the second AUG (AUG2) or a mispaired control PNA (Misp), and analysis by Western blotting using anti-V5 and anti-eGFP antibodies. Results are representative of three independent experiments. D,E C2C12 myoblasts were treated with either the cell-penetrating AUG2 or Misp PNAs, harvested at the indicated time point of a differentiation experiment and subjected to Western blotting using antibodies against TRIP6 / nTRIP6 and GR as a loading control. Representative blots are shown (D). The expression of TRIP6 and nTRIP6 relative to the loading control is presented as mean ± SD of three independent experiments (E). F-J C2C12 cells treated with either the AUG2 or the Misp PNAs were subjected to a differentiation experiment. F,G The relative levels of the *Myog* (F) and *Tnni2* (G) mRNAs were determined by reverse transcription and real-time PCR. Results are plotted relative to the expression of the *Rplp0* gene (mean ± SD of three independent experiments). H-J Cells were fixed at the indicated day and subjected to immunofluorescence analysis using an antibody against MYH3. Nuclei were counterstained using DAPI. H) Representative images are presented (scale bar 200 µm). The percentage of MYH3 expressing mononuclear cells (I) and the fusion index (percentage of nuclei within fused myotubes) (J) are presented as mean ± SD of three independent experiments.

Our previous observation that the levels of nTRIP6 transiently increased during C2C12 myoblast differentiation whereas those of TRIP6 did not vary (Norizadeh Abbariki *et al*, 2021) suggests an increased translation at AUG2. To address this question, we used the AUG2 and the control PNAs fused to a cell-penetrating peptide^15,16^ in a C2C12 differentiation assay. In the control PNA treated cells, nTRIP6 expression transiently increased during the proliferation phase while TRIP6 levels did not vary. The PNA targeting AUG2 abolished the increase in nTRIP6 expression without affecting the levels of TRIP6 (Fig 1D,E). Thus, the transient increase in nTRIP6 expression occurs via increased translation initiation at AUG2. We have previously shown that nTRIP6 represses premature myoblast differentiation, allowing proper myocyte differentiation and fusion at later stages (Norizadeh Abbariki *et al*, 2021). Accordingly, treatment of C2C12 myoblasts with the AUG2 PNA accelerated the expression of *Myog* mRNA, used as an indicator of early myocytic differentiation (Edmondson & Olson, 1989) and delayed the expression of *Tnni2*, a late differentiation gene (Lin *et al*, 1994), as compared to untreated cells or cells treated with the control PNA (Fig 1F,G). Similarly, during the early myocytic differentiation phase (day 1 after the induction of differentiation), the AUG2 PNA reduced the number of cells expressing MYH3 (embryonic myosin heavy chain), another index of late differentiation (Fig 1H,I). Finally, cell fusion, which started at day 2 and strongly increased at day 3 in untreated cells and in cells treated with the control PNA, was significantly inhibited in cells treated with the AUG2 PNA (Fig 1H,J). Thus, the function of nTRIP6 in the temporal regulation of myoblast differentiation and fusion relies on a transient increase in translation at AUG2.

### The mTOR pathway transiently stimulates nTRIP6 translation during myogenesis

In order to identify signaling pathways involved in the upregulation of nTRIP6 translation, we screened a library of 184 unique Medaka kinases (Chen *et al*, 2014; Souren *et al*, 2009) for those altering the nTRIP6/TRIP6 ratio. This ratio was normally distributed (Figure EV2A,B) and analysis of the Z scores (Figure EV2C; Table EV1) revealed 8 kinases which significantly increased it (Table 1), including notably mTOR. Interestingly, AKT1, CHUK and MAP3K7, other hits from the screen, are direct or indirect activators of mTORC1 (Saxton & Sabatini, 2017). We thus tested the effect of mTORC1 on nTRIP6 expression. In proliferating C2C12 myoblasts, treatment with IGF-1, a potent inducer of mTORC1 (Saxton & Sabatini, 2017), increased the expression of nTRIP6 in a dose-dependent manner without affecting that of TRIP6 (Fig 2A,B). Furthermore, a constitutively active mutant of RHEB (RHEB-Q64L) (Jiang & Vogt, 2008), an upstream activator of mTORC1, increased the expression of nTRIP6 from the tagged construct without affecting the expression of TRIP6 (Fig 2C,D). Thus, nTRIP6 translation is stimulated by the mTORC1 pathway. We next investigated whether mTORC1 was responsible for the up-regulation of nTRIP6 translation during C2C12 cell differentiation. mTORC1 activity was elevated in the proliferation phase, reaching a maximum at day -1 (relative to the induction of differentiation), and then decreased prior to differentiation (Fig 2E,F). Treatment with the mTORC1 inhibitor Everolimus abolished the transient increase in nTRIP6 (Fig 2G,H). Thus, the transient increase in mTORC1 activity promotes the increase in nTRIP6 translation during early myogenesis. Interestingly, mTORC1 has been reported to inhibit differentiation at early stages (Ge *et al*, 2011; Wilson *et al*, 2016). Given that nTRIP6 prevents premature differentiation (Norizadeh Abbariki *et al*, 2021) (Fig 1), it is tempting to speculate that mTORC1 inhibits early differentiation at least in part via the up-regulation of nTRIP6 translation.

**Table 1.** Kinases which increase the expression of nTRIP6.

| Kinase | nTRIP6/TRIP6 | Z-score<sup>a</sup> | *P* value |
| --- | --- | --- | --- |
| Mark1 | 0.0964 | 3.016 | 2.56x10<sup>-3</sup> |
| Map3k7 | 0.0896 | 2.578 | 9.94x10<sup>-3</sup> |
| Eif2ak1 | 0.0880 | 2.471 | 1.35x10<sup>-2</sup> |
| Cdc7 | 0.0875 | 2.443 | 1.71x10<sup>-2</sup> |
| Chuk | 0.0866 | 2.385 | 1.71x10<sup>-2</sup> |
| mTor | 0.0847 | 2.258 | 2.39x10<sup>-2</sup> |
| Akt1 | 0.0832 | 2.163 | 3.05x10<sup>-2</sup> |
| Pkn2 | 0.0804 | 1.983 | 4.74x10<sup>-2</sup> |
| Vector<sup>b</sup> | 0.0371±0.015 |  |  |

<sup>a</sup> Z-scores and derived *P* values were determined for the entire kinase set (See Materials and Methods).
<sup>b</sup> The ratio in cells transfected with the empty vector is presented a mean ± SD of 16 determinations.

**Figure 2.**
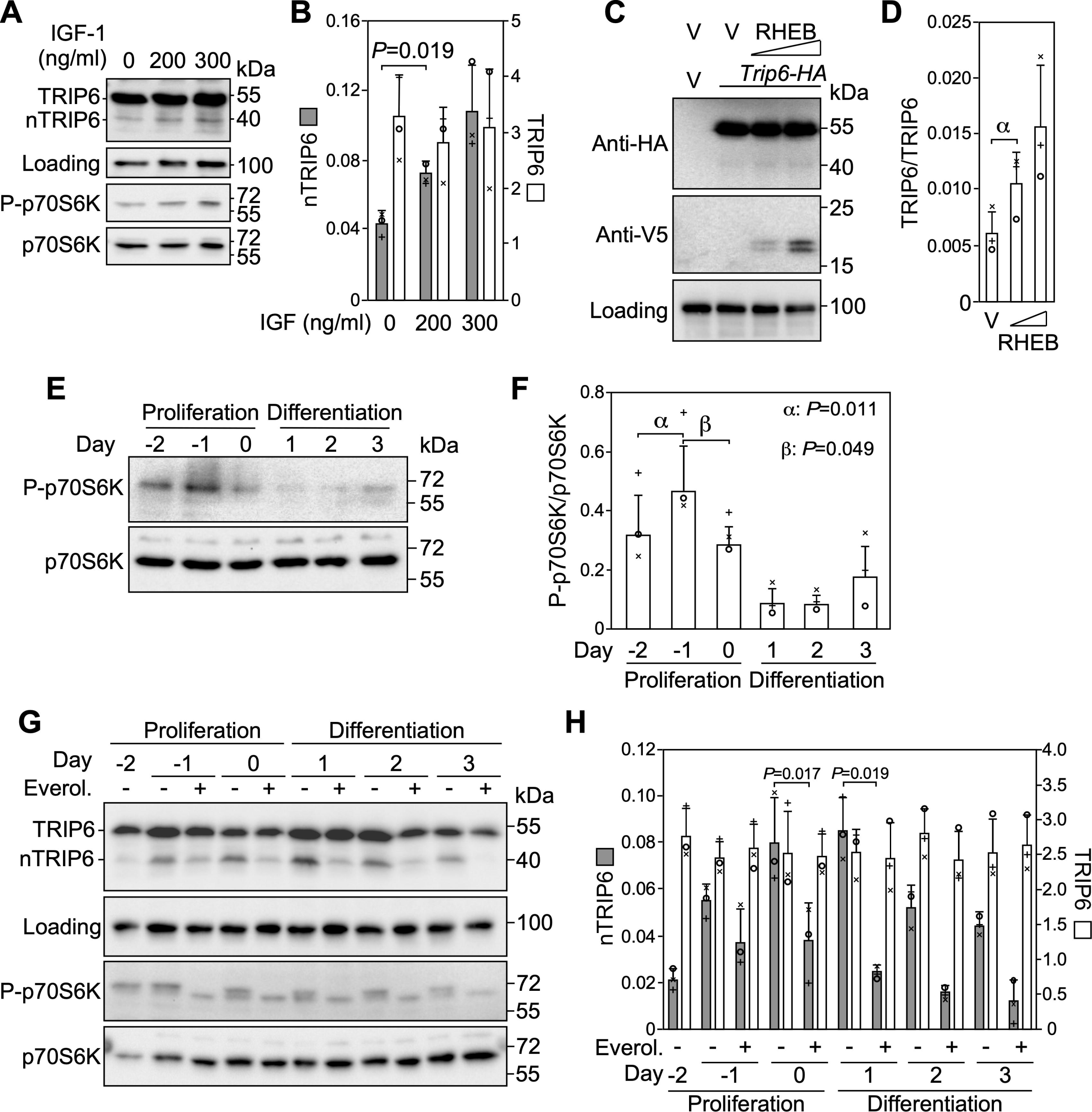
mTORC1 mediates the increase in nTRIP6 translation during myogenesis. A,B Lysates of C2C12 cells treated with IGF-I at the indicated concentration for 6 h were analyzed by Western blotting using antibodies against TRIP6 / nTRIP6, phosphorylated p70S6K (P-p70S6K) and GR as a loading control. Membranes were stripped and re-probed with a phosphorylation-independent p70S6K antibody. Representative Western blots are shown (A). The expression of TRIP6 and nTRIP6 relative to the loading control is presented as mean ± SD of three independent experiments (B). C,D Lysates of C2C12 cells transfected with the HA-tagged *Trip6* construct together with either an empty vector (V) or a V5-tagged constitutively active mutant of RHEB (RHEB-Q64L) at increasing doses were subjected to Western blotting using antibodies against the HA tag, the V5 tag and GR as a loading control. (C) Representative Western blots are shown. (D) The expression of nTRIP6 relative to that of TRIP6 is presented as mean ± SD of three independent experiments (α: *P*=0.032). E,F Lysates of C2C12 myoblasts harvested at the indicated day of a differentiation experiment were subjected to Western blotting using an anti-phosphorylated p70S6 kinase antibody (P-p70S6K). Membranes were stripped and re-probed with a phosphorylation-independent p70S6K antibody. Representative blots are shown (E). The amount of P-p70S6K normalized to the total amount of p70S6K is presented as mean ± SD of three independent experiments (F). G,H) Lysates of C2C12 myoblasts treated as indicated with either solvent alone or Everolimus (Everol.; 100nM) were subjected to Western blotting using antibodies against TRIP6 / nTRIP6, P-p70S6K and GR as a loading control. Membranes were stripped and re-probed with a phosphorylation-independent p70S6K antibody. Representative blots are shown (G). The expression of TRIP6 and nTRIP6 relative to the loading control is presented as mean ± SD of three independent experiments (H). In each graph individual values are depicted by symbols, each representing an independent experiment.

### A short internal upstream open reading frame (iuORF) represses nTRIP6 translation

The stimulatory effect of mTORC1 on nTRIP6 translation is reminiscent of the regulation of translation by short upstream ORFs (uORFs) in the 5’ regulatory regions of mRNAs which typically repress translation initiation at the downstream main ORF translation initiation site (TIS) (Chen & Tarn, 2019; Zhang *et al*, 2019). Indeed, apart from its role in stimulating translation initiation of a subset of mRNAs (Masvidal *et al*, 2017), mTORC1 stimulates re-initiation downstream of uORFs (Calkhoven *et al*, 2000; Zidek *et al*, 2015; Chen *et al*, 2010). Interestingly, the expression of ATF-4, a prototypical protein whose translation is repressed by a uORF (Zhang *et al*, 2019; Chen & Tarn, 2019) was also transiently increased during myoblast differentiation and this increase was also inhibited by Everolimus (Figure EV3). We identified a conserved short ORF in the *Trip6* mRNA coding sequence (Fig 3A; Figure EV4) located immediately upstream of AUG2. Mutation of the short ORF initiation codon into a non-initiating codon in the tagged TRIP6 construct led to an increased expression of nTRIP6 (Fig 3A,B). Furthermore, expression of nTRIP6 from the AUG1-deleted construct was further increased upon mutation of the short ORF AUG (Fig 3A,B), confirming the repressive role of the short ORF for initiation at AUG2 after leaky scanning through AUG1. Although ribosome profiling experiments have revealed the existence of short ORFs in the coding sequences of mRNAs (Ingolia *et al*, 2011; Bazzini *et al*, 2014), their function remained elusive. Our results show that the short ORF within *Trip6* coding sequence behaves like “classical” repressive uORFs in 5’ leader sequences. We therefore designated this regulatory element an internal uORF (iuORF).

**Figure 3.**
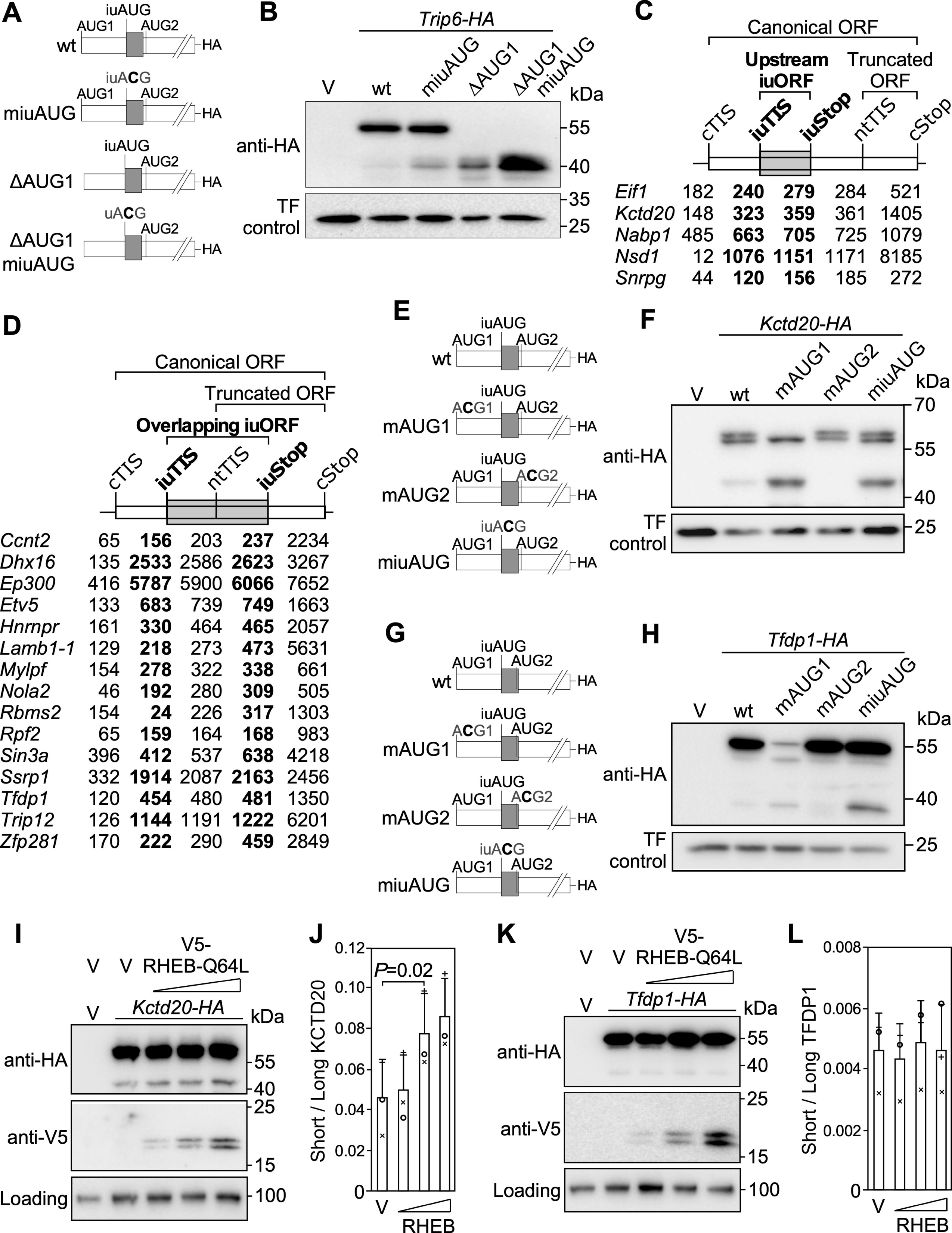
Short internal ORFs repress the translation on N-terminally truncated proteoforms. A Schematic representation of the TRIP6 constructs used. The internal upstream open reading frame (iuORF) is indicated as a grey box. B Lysates of NIH-3T3 fibroblasts transfected with either an empty vector (V) or the indicated constructs, together with eGFP as a transfection (TF) control were analyzed by Western blotting using anti-HA and anti-eGFP antibodies. Results are representative of three independent experiments. C,D Schematic representation of other mRNAs harboring upstream (C) or overlapping (D) iuORFs. cTIS: canonical translation initiation site; iuTIS and iuSTOP: TIS and stop codon for the iuORF; nTIS: TIS for the N-terminally truncated proteoform ORF; cSTOP: canonical stop codon. The numbers indicate the positions on the mRNA. E-H NIH-3T3 fibroblasts were transfected with either an empty vector (V) or the indicated C-terminally tagged *Kctd20* (E,F) or *Tfdp1* (G,H) CDS construct together with mCherry as a transfection (TF) control. Cell lysates were analyzed by Western blotting using anti-HA and anti-mCherry antibodies. Results are representative of three independent experiments. I-L Lysates of C2C12 cells transfected with either *Kctd20-HA* (I,J) or *Tfdp1-HA* (K,L) together with either an empty vector (V) or the V5-tagged constitutively active mutant RHEB-Q64L at increasing doses were subjected to Western blotting using antibodies against the HA tag, the V5 tag and GR as a loading control. (I,K) Representative Western blots are shown. The expression of the short proteoform relative to that of the long proteoform of KCTD20 (J) or TFDP1 (L) is presented as mean ± SD of three independent experiments; individual values are depicted by symbols, each representing an independent experiment.

To investigate whether other mRNAs also harbor iuORFs, we curated a published ribosome profiling dataset that identified over 13,000 TISs in mouse embryonic stem cells (Ingolia *et al*, 2011). Based on stringent filtration criteria (see Materials and Methods section), among the 4,994 genes analyzed we identified 20 mRNAs exhibiting a short internal ORF in another frame than the canonical ORF and in close proximity to a TIS for an N-terminally truncated proteoform. Among these short internal ORFs 5 were upstream, with the stop codon immediately upstream of the truncated proteoform TIS (Fig 3C), and 15 were overlapping, with the stop codon downstream of the truncated proteoform TIS (Fig 3D). In the case of uORFs in 5’ regulatory regions, the sequence context of the initiation codon, the length of the short ORF and its distance to the downstream TIS are the most relevant features to regulate initiation at the next downstream TIS (Chen & Tarn, 2019; Zhang *et al*, 2019). In the identified short internal ORFs these features were highly conserved in the human and rat mRNAs (Table EV2). We validated this finding by studying the contribution of the identified TISs in *Kctd20* mRNA, which bears an upstream iuORF and in *Tfdp1* mRNA which harbors an overlapping iuORF (Fig 3E-H). Transfection of NIH-3T3 cells with a C-terminally tagged KCTD20 construct gave rise to a protein with a size corresponding to the usage of the annotated first AUG, which was indeed not expressed when this AUG was mutated into a non-initiating codon. A slightly smaller protein, presumably translated from a downstream non-canonical TIS was also detected, which was still present when AUG1 was mutated. The wild-type construct also gave rise to a short proteoform with a size corresponding to the predicted size of the postulated N-terminally truncated proteoform. Its expression was strongly increased when the first AUG was mutated, and was abolished when the AUG for the postulated N-terminally truncated proteoform (AUG2) was mutated into a non-initiating codon (Fig 3E,F). These results confirm that this short proteoform is translated from AUG2 after leaky scanning at AUG1. Furthermore, mutation of the identified upstream iuORF initiation codon into a non-initiating codon led to an increased expression of the short proteoform (Fig 3E,F). Similarly, a C-terminally tagged TFDP1 construct gave rise to a protein with a size corresponding to the usage of the annotated first AUG, which was not expressed when this AUG was mutated into a non-initiating codon. A shorter proteoform with a size corresponding to that of the postulated N-terminally truncated proteoform was expressed at very low levels from the wild-type construct. Its expression was slightly increased from the construct with a mutated AUG1, and strongly increased when the identified overlapping iuORF initiation codon was mutated into a non-initiating codon (Fig 3G,H).

Furthermore, similarly to what we observed with nTRIP6, activation of mTORC1 by transfection of RHEB-Q64L increased the expression of the short KCTD20 proteoform from the tagged construct without affecting the expression of canonical long proteoform (Fig 3I,J). Thus, the translation of N-terminally truncated proteoforms can be regulated independently of the translation of the canonical ORF. However, neither the short nor the canonical long proteoforms of TFDP1 were regulated by RHEB-Q64L (Fig 3K,L). mTORC1 has been shown to inhibit post-termination ribosome recycling after the translation of uORFs, and thereby to promote re-initiation at downstream start codons (Schepetilnikov *et al*, 2013; Gunišová *et al*, 2018; Zidek *et al*, 2015). Thus, mTORC1 most likely favors re-initiation at the nTRIP6 and at the short KCTD20 proteoform AUGs after translation of the iuORF. The iuORF in *Tfdp1* mRNA overlaps with the TIS for the short proteoform. Consequently, re-initiation downstream of the iuORF cannot occur at this TIS. Therefore, it seems very likely that mTORC1 stimulates re-initiation downstream of upstream iuORFs to increase the translation of the N-terminally truncated proteoforms.

In conclusion, we have characterized iuORFs as novel mRNA cis-regulatory elements which regulate the translation of N-terminally truncated proteoforms.

## Materials and Methods

### Plasmid constructs

In pcDNA3.1-TRIP6-HA the mouse *Trip6* CDS with a C-terminal HA epitope was cloned by PCR between the NheI and EcoRI sites of pcDNA3.1 (Invitrogen, Karlsruhe, Germany). The KpnI / XhoI fragment of pcDNA3.1-TRIP6-HA, lacking the first ATG, was cloned into pcDNA3.1 to obtain pcDNA3.1-TRIP6ΔAUG1-HA. pcDNA3.1-TRIP6-mAUG2-HA and pcDNA3.1-TRIP6-iuACG-HA were obtained by mutating the nTRIP6 ATG (ATG2) and the iuORF ATG (iuATG) into ACG, respectively, using the QuikChange site-directed mutagenesis kit (Agilent, Waldbronn, Germany). The KpnI / XhoI fragment of pcDNA3.1-TRIP6-iuACG-HA was cloned into pcDNA3.1 to obtain pcDNA3.1-TRIP6ΔAUG1-iuACG-HA. The *Trip6* CDS constructs C-terminally tagged with the V5 epitope pcDNA3.1-TRIP6-V5 and pcDNA3.1-TRIP6ΔAUG1-V5 were cloned using a similar strategy. The constitutively active RHEB mutant RHEB-Q64L (Jiang & Vogt, 2008) N-terminally tagged with the V5 epitope was synthesized as a DNA string (GeneArt Strings DNA Fragments, Thermo Fisher Scientific, Schwerte, Germany) and cloned into pcDNA3.1 to obtain pcDNA3.1-V5-RHEB-Q64L. C-terminally HA-tagged KCTD20 and TFDP1 constructs were cloned as synthesized DNA strings into pcDNA3.1. The QuikChange site-directed mutagenesis kit was used to mutate the canonical initiation codon (AUG1), the initiation codon for the N-terminally truncated proteoform (AUG2) and the initiation codon for the short internal ORF (iuAUG) into non-initiating codons (ACG) to obtain pcDNA3.1-KCTD20-mAUG1-HA, pcDNA3.1-KCTD20-mAUG2-HA, pcDNA3.1-KCTD20-miuAUG-HA, pcDNA3.1-TFDP1-mAUG1-HA, pcDNA3.1-TFDP1-mAUG2-HA and pcDNA3.1-TFDP1-miuAUG-HA. pcDNA3.1-eGFP and pcDNA3.1-mCherry were obtained by cloning the corresponding string DNA fragments (ThermoFisher Scientific) into pcDNA3.1. All constructs were verified by sequencing.

### Cell culture, transfection and cellular assays

C2C12 myoblasts, NIH-3T3 fibroblasts and HEK-293 cells (all from ATCC, LGC Standards GmbH, Wesel, Germany) were cultured in Dulbecco’s modified Eagle’s medium (DMEM) supplemented with 10% fetal calf serum (FCS). Cell lines were authenticated by morphological examination and were routinely checked for mycoplasma contamination.

We used a standardized protocol for the differentiation of C2C12 myoblasts (Norizadeh Abbariki *et al*, 2021). Cells were seeded at a density of 5×10^3^ cells / cm^2^ in growth medium (GM, 10% FCS-containing DMEM) at day -3, relative to the induction of differentiation at day 0. When cells reached at day 0, differentiation was induced by changing the medium to differentiation medium (DM, 2% horse serum-containing DMEM), which was then replaced every second day. Where indicated, cells were treated with the mTORC1 inhibitor Everolimus (Sigma-Aldrich) at a concentration of 100 nM. NIH-3T3 fibroblasts and C2C12 myoblasts were transfected using *Trans*IT-X2 (Mirus Bio LLC) and HEK-293 cells with ScreenFect-A (ScreenFect, Eggenstein-Leopoldshafen, Germany). A PNA was designed to target the mouse *Trip6* mRNA sequence (NM_011639.3) from positions 481 to 498 which encompass AUG2: NH_2_-GTCCAGATCAGCCAACAT-R_8_-CONH_2_. A mispaired PNA (Misp) was used as a control: NH_2_-<u>C</u>TCCA<u>C</u>ATCA<u>C</u>CCAAGAT-R_8_-CONH_2_; the mispaired bases are underlined. The cell-penetrating moiety of the PNAs consists of an octoarginine peptide (R8). The PNAs were synthesized manually by standard Fmoc-based PNA solid-phase protocols as described (Vázquez & Seitz, 2014) (TentaGel® S RAM resin; 10 μmol scale; after the 12^th^ PNA-monomer coupling, the used equivalents increased from 4 to 8 for 1h). Peptide-PNA conjugates were purified by preparative reverse-phase (RP) HPLC and identified by high-resolution mass spectrometry (Figure EV1). To increase the penetration, the PNAs were applied to C2C12 cells at a concentration of 10 µM in DMEM with reduced FCS (2.5%) for 2h, after which FCS concentration was increased to 10%.

For the kinase over-expression screen, a library of 184 unique Medaka kinases cloned into pCMV-Sport6.1 (Chen *et al*, 2014; Souren *et al*, 2009) was kindly provided by Jochen Wittbrodt (Centre for Organismal Studies, Heidelberg, Germany) and Gary Davidson (Institute for Biological and Chemical Systems, Karlsruhe Institute of Technology, Karlsruhe, Germany). HEK-293 cells were co-transfected in 96-well plates with the V5-tagged TRIP6 construct and either an empty vector as a control or the library using ScreenFect-A. Cells were lysed 24h later, and the relative expression of TRIP6 and nTRIP6 was assessed by Western Blot analysis (see below).

### RNA isolation and quantitative Real-time PCR (qRT-PCR)

Total RNA was extracted using PeqGOLD TriFast^TM^ (Peqlab Biotechnologie, Erlangen, Germany) and reverse-transcribed into cDNA. *Myog* (myogenin) and *Tnni2* mRNAs, as well as the transcript of the large ribosomal subunit P0 gene (*Rplp0*) used for normalization, were quantified by real-time PCR using the ABI Prism Sequence Detection System 7000 (Applied Biosystems, Foster City, CA). The primers (ThermoFisher Scientific) were as follows (5’ to 3’): Myog: GAGACATCCCCCTATTTCTACCA and GCTCAGTCCGCTCATAGCC; Tnni2: CATGGAGGTGAAGGTGCAGA and CTCTTGAACTTGCCCCTCAGG; *Rplp0*: GGACCCGAGAAGACCTCCTT and GCACATCACTCAGAATTTCAATGG.

### Western Blotting

Western blotting analyses were performed using the following antibodies: a custom-made rabbit anti-TRIP6 monoclonal antibody (Kemler *et al*, 2016); anti-V5 (R960-25, Thermo Fisher Scientific); anti-HA (clone 3F10, Roche Applied Science, Mannheim, Germany); anti-GFP (clone D5.1, Cell Signaling Technology); anti-mCherry (Ab167453, Abcam); anti-ATF4 (11815, Cell Signaling Technology); anti-phospho (Thr389)-p70 S6 Kinase (clone 108D2, Cell Signaling Technology). To normalize the phospho (Thr389)-p70 S6 Kinase signals membranes were stripped and reprobed with a phosphorylation-independent anti-p70 S6 Kinase antibody (clone 49D7, Cell Signaling Technology). The expression of the “classical” housekeeping genes β-actin and GAPDH is regulated during myogenesis (Hildyard & Wells, 2014; Otey *et al*, 1988; Cox *et al*, 1990). Therefore, we selected the glucocorticoid receptor (GR) as a loading control, given that it is stably expressed during C2C12 differentiation (Sun *et al*, 2008). To this end we used an anti-GR antibody (sc-393232, Santa Cruz, Heidelberg, Germany). Signals were detected by enhanced chemoluminescence using the ChemiDoc Touch Imaging System (BioRad laboratories, Munich, Germany). Signal quantification was performed within the linear range of detection using the Image Lab software (Bio-Rad laboratories). Linear brightness and contrast adjustments were made for illustration purposes only after the analysis had been made.

### *In vitro* transcription and translation

The plasmids pcDNA3.1-TRIP6-V5 and pcDNA3.1-TRIP6ΔAUG1-V5, as well as pcDNA3.1-eGFP as a control were used as templates for *in vitro* transcription and translation using the T7 TNT^®^ Coupled Reticulocyte Lysate System (Promega, Mannheim, Germany) according to the manufacturer’s instructions, except that the reactions were performed for 90 min at 30°C in the presence of 5 μM PNA targeting the 2^nd^ AUG of *Trip6* mRNA or its mispaired control. The translated products were analyzed by Western Blotting.

### *In silico* identification of iuORFs

In order to identify other mRNAs harboring short internal ORFs that putatively regulate the translation of N-terminally truncated proteoforms, we curated a published ribosome profiling dataset (Ingolia *et al*, 2011). Out of the 4,994 mRNAs analyzed, we first selected those in which the canonical TIS was identified and that harbored at least one internal (3’ to the canonical TIS) out of frame, short (≤ 100 codons) ORF and at least one in frame internal TIS for a truncated proteoform. The resulting mRNAs were then filtered to keep only those in which the TIS of the short ORF was located 5’ to the truncated proteoform TIS, and in which the stop codon of the short ORF was located either 3’ to the truncated proteoform TIS (overlapping short ORF) or at a maximum distance of 30 nucleotide 5’ to the truncated proteoform TIS (upstream short ORF). When several short ORFs fulfilled these criteria within the same mRNA, only the one with the highest initiation score (Ingolia *et al*, 2011) was kept. We then performed a conservation analysis of the features most relevant for the translation of N-terminally truncated proteoforms and for short ORFs to regulate initiation at downstream AUGs (Chen & Tarn, 2019; Zhang *et al*, 2019), i.e. the sequence contexts of the canonical, short ORF and truncated proteoform initiation codons relative to the Kozak sequence (Kozak, 2002), the distance from the canonical TIS to the short ORF TIS, the distance from the short ORF stop codon to the truncated proteoform TIS and the length of the short ORF. Human and rat orthologues to mouse genes were selected using the closest homologue in the phylogenetic tree using clustalw. Conservation of the start codons and length of the ORF were obtained by screening the multiple alignments with the EMBOSS (Rice *et al*, 2000) functions extractalign and transeq.

### Immunofluorescence, microscopy and image analysis

Immunofluorescence analysis was performed on C2C12 cells grown and differentiated on glass coverslips coated with collagen Type I (Sigma-Aldrich), fixed for 10 min in 10% formalin, permeabilized for 10 min in 0.5% Triton X-100 in PBS and blocked for 1h in 5% BSA in PBS. The primary antibody was a mouse anti-MYH3 (F1.652-b, Developmental Studies Hybridoma Bank, deposited by Blau H.M.) and the secondary antibody was an Alexa Fluor 488-conjugated anti-mouse antibody (Invitrogen). Nuclei were counter-stained with DAPI (Thermo Fisher Scientific), Cells were imaged by confocal microscopy on a Zeiss LSM 800 (Zeiss, Jena, Germany). Cells images were acquired in tiling mode using a 10x/0.3 Plan-Neofluar objective resulting in 3 X 2 mm^2^ images. Images were analyzed using Fiji (Schindelin *et al*, 2012). The number of MYH3 positive mononuclear cells was determined by a combination of automated segmentation and manual counting. The fusion index was calculated as the percentage of nuclei within fused myotubes. Linear brightness and contrast adjustments were made for illustration purposes, but only after the analysis had been made.

### Statistical analysis

Statistical analyses were performed using R. Where indicated, significant differences were assessed by two-sided t-test analysis, with values of *P* < 0.05 sufficient to reject the null hypothesis. A Bonferroni correction was applied when multiple comparisons were performed. For the kinase overexpression screen, the normal distribution of the nTRIP6/TRIP6 ratio was first confirmed by a Shapiro-Wilk normality test (W = 0.9803, *P* = 1.07×10^-2^) as well as by the Kernel density and Q-Q plots (Figure EV2). This allowed us to calculate Z scores and to derive 2-sided *P* values in order to determine the “hits”, kinases which significantly altered the nTRIP6/TRIP6 ratio.

## Acknowledgments

The authors are grateful to Jochen Wittbrodt (Centre for Organismal Studies, Heidelberg, Germany) and Gary Davidson (Institute for Biological and Chemical Systems, Karlsruhe Institute of Technology, Karlsruhe, Germany) for providing the Medaka kinase library. We thank Nicholas S. Foulkes and Andrew C.B. Cato for stimulating discussions and critically reading the manuscript. This work was supported in part by the German Science Foundation (grants 315384510 to O.K. and 425970020 to O.K. and O.V.). We acknowledge support by the KIT-Publication Fund of the Karlsruhe Institute of Technology.

## Author contributions

Conceptualization, O.K. and O.V.; Methodology, R.F., Z.G., C.T., M.W., M.L., O.A., O.V. and O.K.; Investigation, R.F, Z.G., N.W., P.S., C.T., M.W., S.B., M.L. and C.E.; Writing – Original Draft, O.K.; Writing – Review & Editing, C.E., O.A., O.V. and O.K.; Funding Acquisition, O.V. and O.K.; Resources, L.Z., G.L., M.L. and C.E.; Supervision, O.V. and O.K.

## Data availability

This study includes no data deposited in external repositories.

## Expanded view Figure legends

**Figure EV1. HPLC and high-resolution mass spectrometry analysis of the cell-penetrating AUG2 targeting (A) and mismatched control (B) peptide nucleic acids.**

**Figure EV2. Relative expression of nTRIP6 and TRIP6 in the kinase over-expression screen.** The nTRIP6/TRIP6 ratios were determined by Western blotting analysis in HEK-293 co-transfected with the C-terminally V5-tagged Trip6 CDS construct and a library of 184 unique Medaka kinases. A,B) The ratios are normally distributed. The Kernel density and Q-Q plots are presented. Shapiro-Wilk normality test: W = 0.9803, P = 1.07×10-2. C) Z scores of the nTRIP6/TRIP6 ratios.

**Figure EV3. mTORC1-dependent regulation of ATF4 expression during myogenesis.**

C2C12 myoblasts were treated as indicated with either solvent alone or Everolimus (100nM) and harvested at the indicated time point of a differentiation experiment. Cell lysates were subjected to Western blotting using antibodies against ATF4 and GR as a loading control. A) Representative blots are shown. B) The expression of ATF4 relative to the loading control is presented as mean ± SD of three independent experiments.

**Figure EV4. Conservation of the internal upstream open reading frame (iuORF) in Trip6 mRNA.**

For each species, the schematic representation of the 5’ of Trip6 mRNA depicts the surrounding sequence of the TRIP6 (AUG1), iuORF and nTRIP6 (AUG2) initiation codons relative to the Kozak sequence (RNN AUG GNN where N represents any nucleotide and R a purine), the number of nucleotides (nt) in the uORF and the distance between the iuORF and AUG2. The grey box represents the Nuclear Export Signal encoding sequence.

