## Supplementary figures and images for "Short internal open reading frames regulate the translation of N-terminally truncated proteoforms"

### Supplementary figure 1

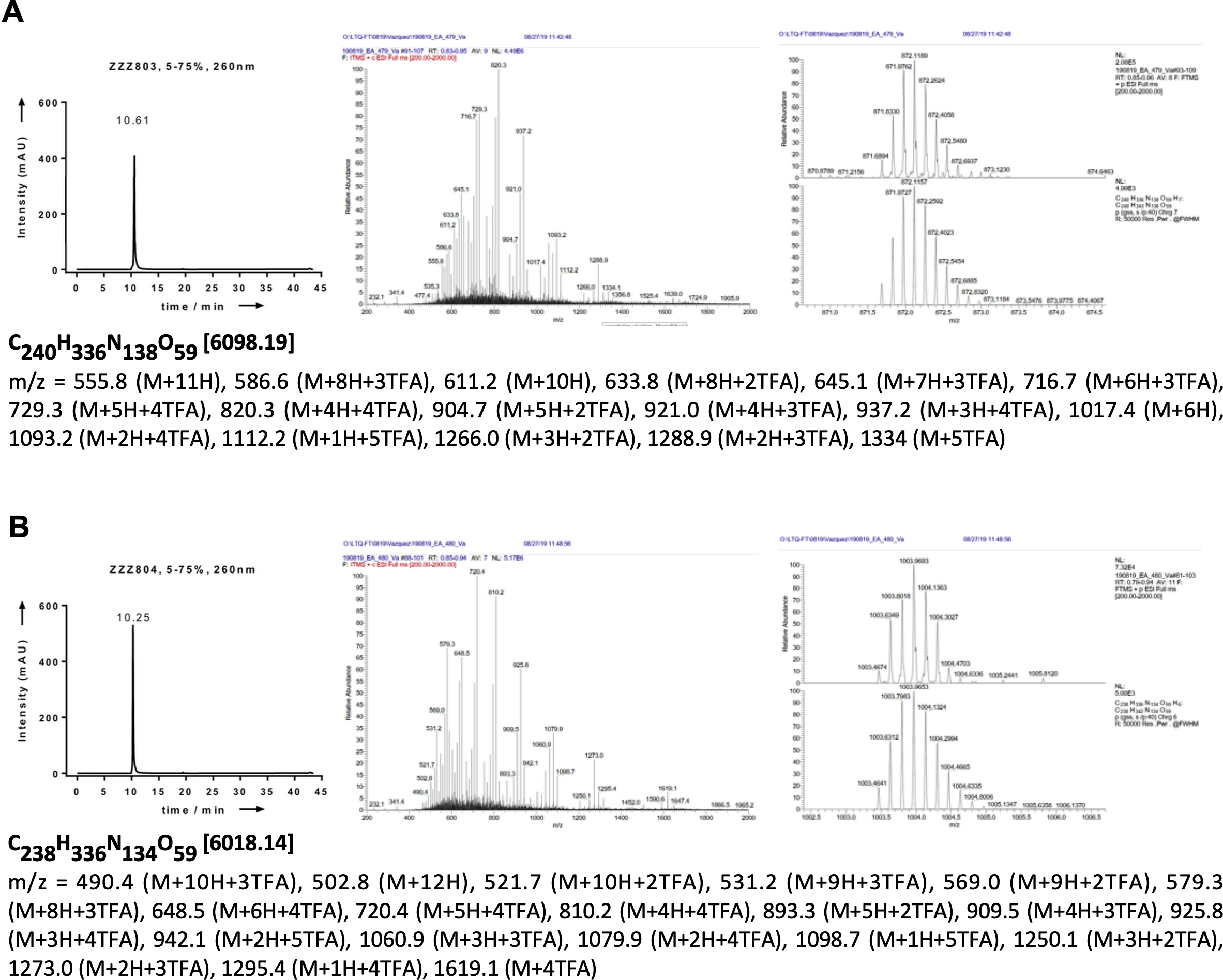

### Supplementary figure 2

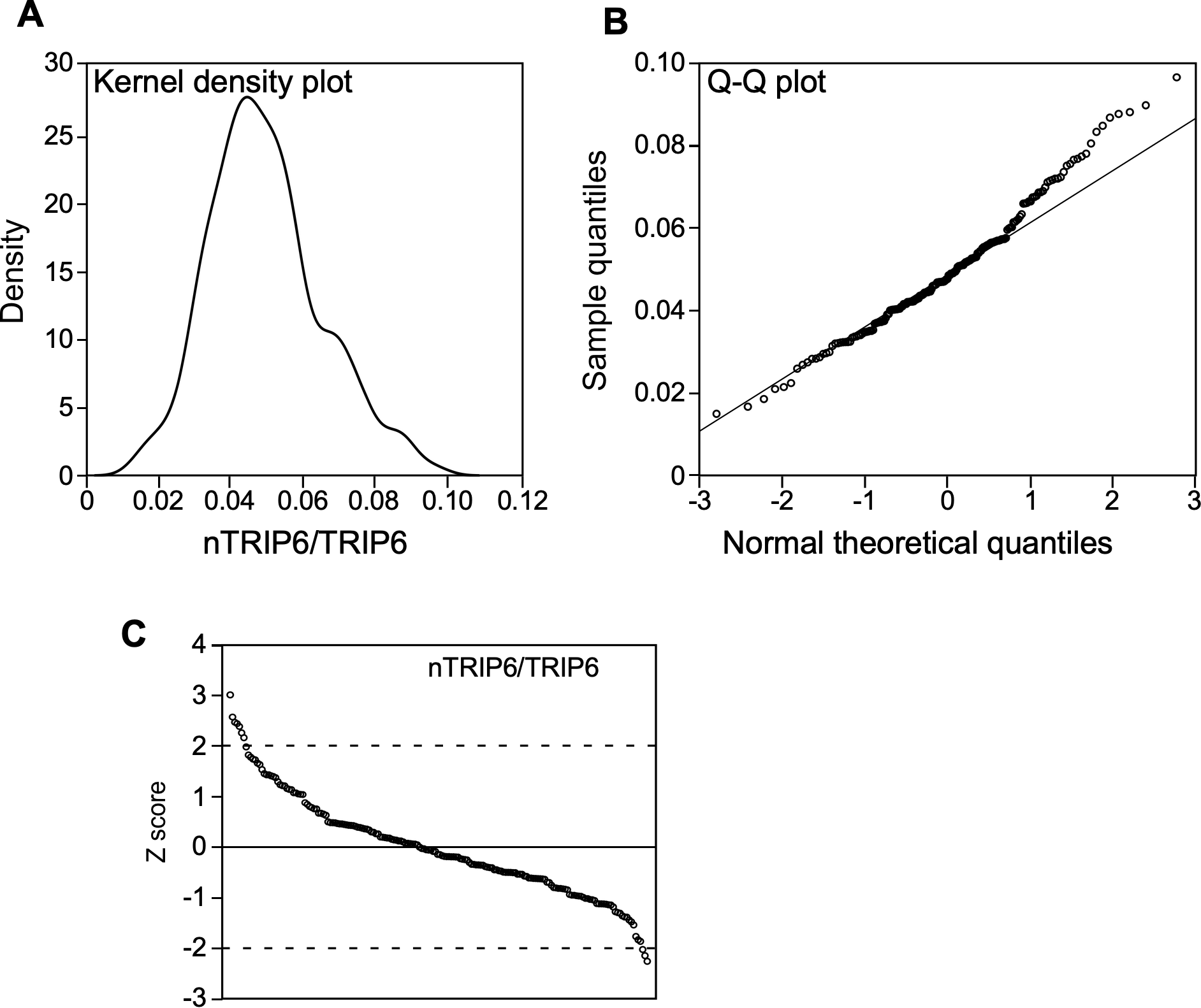

### Supplementary figure 3

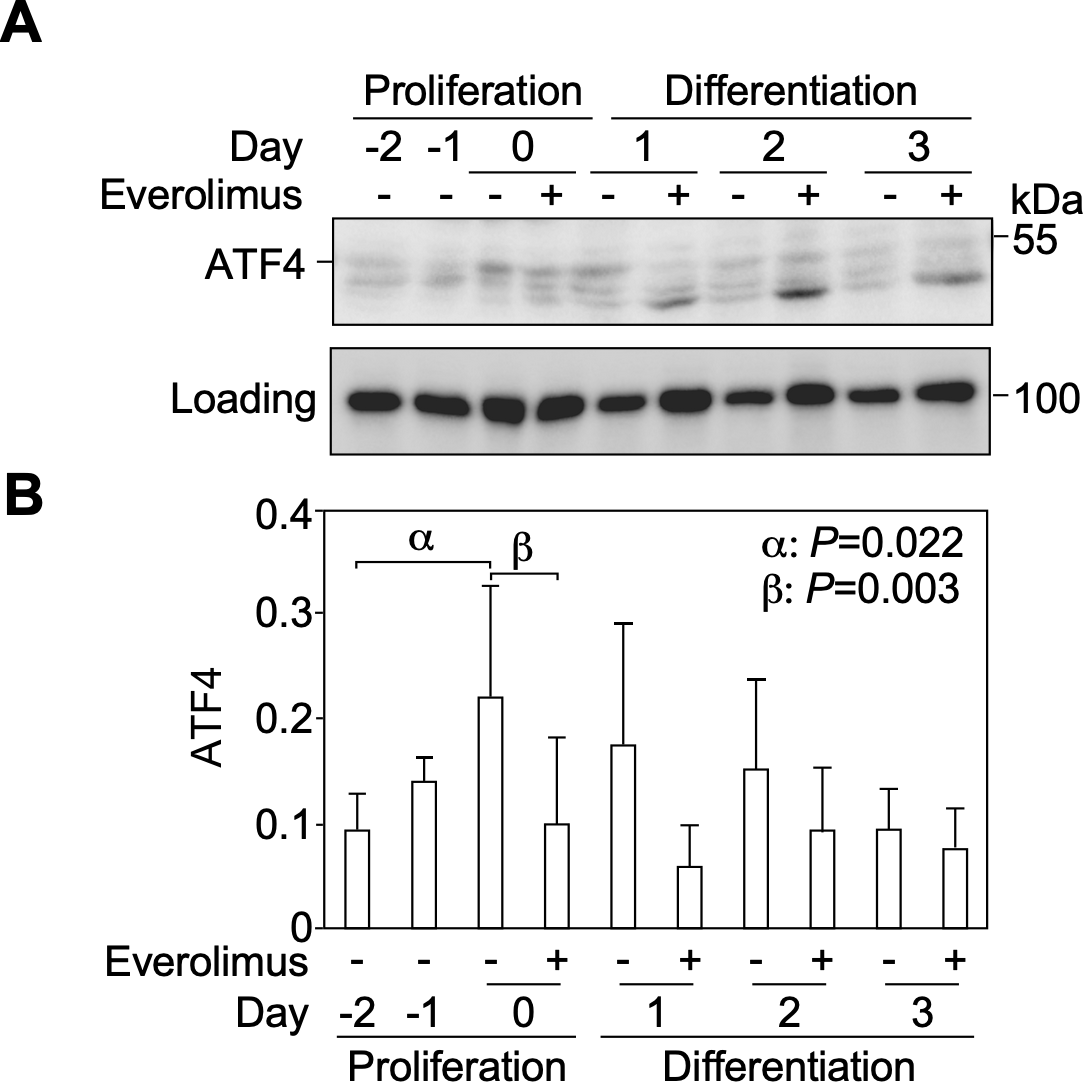

### Supplementary figure 4

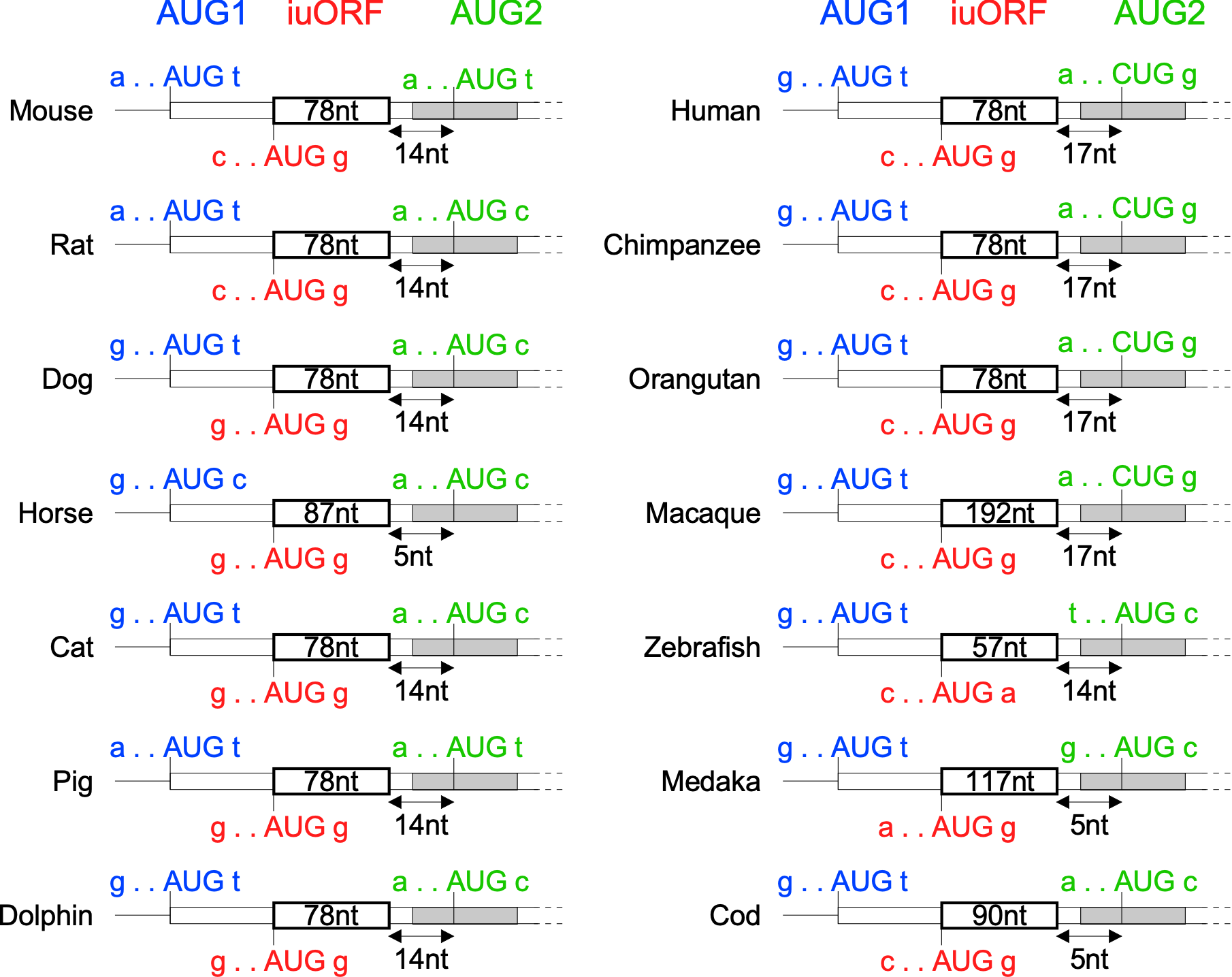
