## Supplementary table 1 for "Short internal open reading frames regulate the translation of N-terminally truncated proteoforms": README_Table EV1.docx

**Table EV1. Effect of overexpressed kinases on the relative expression of nTRIP6 and TRIP6.**

HEK-293 cells were co-transfected with the V5-tagged TRIP6 construct and either an empty vector as a control or a library of 184 unique Medaka kinases. Cells were lysed 24h later, and the relative expression of TRIP6 and nTRIP6 was assessed by Western Blot analysis. For each kinase, the nTRIP6/TRIP6 ratio, the Z score and the *P* value is indicated.
