## Supplementary table 2 for "Short internal open reading frames regulate the translation of N-terminally truncated proteoforms": README_Table EV2.docx

**Table EV2. Conservation of the short internal ORFs and translation initiation sites for N-terminally truncated proteoforms.**

For each identified mRNA exhibiting a short internal ORF in close proximity to a translation initiation site (TIS) for an N-terminally truncated proteoform, the table presents the conservation between the mouse, rat and human sequences of the features most relevant for the translation of N-terminally truncated proteoforms and for short ORFs to regulate initiation at downstream codons.
